# Corynebacterium of the *diphtheriae* complex in companion animals: clinical and microbiological characterization of 64 cases from France

**DOI:** 10.1101/2023.01.04.522820

**Authors:** Kristina Museux, Gabriele Arcari, Guido Rodrigo, Melanie Hennart, Edgar Badell, Julie Toubiana, Sylvain Brisse

## Abstract

**Objectives:** Corynebacteria of the *diphtheriae* complex (Cdc) can cause diphtheria in humans and have been reported from companion animals. We aimed to describe animal infection cases caused by *Cdc* isolates.

**Methods:** 18 308 animals (dogs, cats, horses and small mammals) with rhinitis, dermatitis, non-healing wounds and otitis were sampled in metropolitan France (August 2019 to August 2021). Data on symptoms, age, breed, and the administrative region of origin were collected. Cultured bacteria were analyzed for *tox* gene presence, for production of the diphtheria toxin, for antimicrobial susceptibility, and genotyped by multilocus sequence typing.

**Results:** *C. ulcerans* was identified in 51 cases, 24 of which were toxigenic. Rhinitis was the most frequent presentation (18/51). Eleven cases (6 cats, 4 dogs, 1 rat) were mono-infections. Large breed dogs, especially German Shepherds (9 of 28 dogs; p < 0.00001) were overrepresented. *C. ulcerans* isolates were susceptible to all tested antibiotics. *tox-*positive *C. diphtheriae* was identified in 2 horses. Last, 11 infections cases (9 dogs, 2 cats; mostly chronic otitis, and 2 sores) had *tox*-negative *C. rouxii*, a recently defined species. *C. rouxii* and *C. diphtheriae* isolates were susceptible to most antibiotics tested, and almost all of these infections were polymicrobial.

**Conclusions:** Monoinfections with *C. ulcerans* point towards a primary pathogenic potential to animals. *C. ulcerans* represents an important zoonotic risk, and *C. rouxii* may represent a novel zoonotic agent. This case series provides novel clinical and microbiological data on *Cdc* infections, and underlines the need for management of animals and their human contacts.

**Importance:** We report on the occurrence, clinical and microbiological characteristics of infections caused by members of the Corynebacteria of the *diphtheriae* complex (Cdc) in companion animals. This is the first study based on the systematic analysis of a very large animal cohort (18 308 samples), which provides data on the frequency of Cdc isolates in various types of clinical samples from animals. Awareness of this zoonotic bacterial group remains low among veterinarians and veterinary laboratories, among which it is often considered a commensal bacteria of animals. We suggest that in case of Cdc detection in animals, the veterinary laboratories should be encouraged to send the samples to a reference laboratory for analysis of the presence of the *tox* gene. This work is relevant to the development of guidelines in case of Cdc infections in animals, and underlines their public health relevance given the zoonotic transmission risk.

## Introduction

Diphtheria is a potentially fatal infection in humans, caused mostly by toxigenic *Corynebacterium (C*.*) diphtheriae* isolates, which carry the *tox* gene coding for diphtheria toxin. This bacterial species is phylogenetically related to 5 other *Corynebacterium* species and together with these, is grouped into the *C. diphtheriae* complex (Cdc). *C. ulcerans* (Riegel et al., 1995) can be isolated from humans and animals and is being increasingly reported (Hacker et al., 2016a; Robert Koch-Institut, 2018; Haut Conseil de la santé publique, 2019; Wagner et al., 2010). *C. pseudotuberculosis* causes caseous lymphadenitis in small ruminants and oedomatous skin disease in buffaloes. Although this species is considered potentially toxigenic, only isolates from buffalos in Egypt were reported to produce diphtheria toxin, and there is no evidence for toxigenicity of recent isolates from caseous lymphadenitis (Schlicher et al., 2021; Selim, 2001; Selim et al., 2016; Viana et al., 2017). Two novel species, *C. belfantii* and *C. rouxii*, were recenty described (Badell et al., 2020; Dazas et al., 2018). Isolates of these species were previously identified as *C. diphtheriae*, are mostly *tox*-negative, and are of biovar Belfanti (starch and nitrate negative). Last, *C. silvaticum* was recently described from wild boars; all isolates of this species carry a disrupted *tox* gene, impairing their capacity to produce diphtheria toxin (Dangel et al., 2020). Hence, although all members of the Cdc may potentially harbor the *tox* gene, toxigenic strains are frequently encountered only in *C. diphtheriae* and *C. ulcerans*. In *C. diphtheriae*, this gene is carried on a temperate phage that has integrated into the chromosome of a large variety of sublineages (Guglielmini et al., 2021; Hennart et al., 2020). In *C. ulcerans*, in addition to a lysogenic phage, the *tox* gene can be carried on a pathogenicity island (Dangel et al., 2019). Strains that carry the *tox* gene generally produce the diphtheria toxin *in vitro*, but a short fraction of them (∼10-15%) do not, due to disruptions of the *tox* gene; these are called non-toxigenic, *tox*-bearing (NTTB) strains and are observed both in *C. diphtheriae* and in *C. ulcerans* (and in all *C. silvaticum*).

Although classically defined as a respiratory infection caused by toxigenic *C. diphtheriae*, diphtheria is sometimes defined more broadly as any infection (respiratory, cutaneous or else) potentially leading to manifestations due to the production of the diphtheria toxin by species of the Cdc (https://www.ecdc.europa.eu/en/diphtheria). Following recent taxonomic updates, diphtheria may be defined more broadly as any infection caused by any isolate of the Cdc, irrespective of toxigenic status (Haut Conseil de la santé publique, 2021).

The typical clinical expressions of diphtheria in human are: (i) classical respiratory diphtheria with pseudo-membranous angina, that can provoke a deadly obstruction of upper airways, usually associated with fever and enlarged anterior cervical lymph nodes and oedema (‘bull neck’ appearance); (ii) Cutaneous manifestations, with “rolled edge” ulcers usually observed on the limbs and that can be covered by a greyish pseudo-membrane; and (iii) Toxigenic complications such as polyneuropathy or myocarditis (Sharma et al., 2019; UK Health Security Agency, 2022). Seroprevalence studies in humans demonstrated that up to 82% of the population has titers of anti-diphtheria toxin antibodies below the limit of protection, and that the proportion of non-protected persons increases with age (Berbers et al., 2021; Launay et al., 2009).

Before 1999, no case of *C. ulcerans* were reported in France (Hommez et al., 1999); but, between 2002 and 2013, 28 cases of toxigenic *C. ulcerans* have been reported and between January 2018 and August 2019, 11 further human clinical cases were described; most of the cases were indigenous (Haut Conseil de la santé publique, 2019; National_Reference_Center_for_diphtheria_in_France, 2021) and a similar pattern of increased reporting of *C. ulcerans* was reported from the UK and Germany (European Centre for Disease Prevention and Control, 2021; Hacker et al., 2016a; Robert Koch-Institut, 2018; Wagner et al., 2010).

While the transmission of *C. diphtheriae* is essentially inter-human, *C. ulcerans* is a zoonotic pathogen (Hacker et al., 2016a; Othieno et al., 2019). No transmission of *C. ulcerans* among humans was reported since its description in 1995; however, the possiblity of person-to-person transmission cannot be totally excluded (Konrad et al., 2015). Animals from which *C. ulcerans* is isolated can be asymptomatic carriers but can also present clinical symptoms, such as ulcerative dermatitis and chronic rhinitis (Lartigue et al., 2005; Carfora et al., 2018; Katsukawa et al., 2012; Abbott et al., 2020). Horses can also carry *C. ulcerans* and sometimes show signs of respiratory diphtheria (Zendri et al., 2021).

Similarly, *C. rouxii* may also be zoonotic: no interhuman transmission has been reported yet, and in addition to human cases, it has been so far identified in dogs, cats and a fox (Badell et al., 2020; Hall et al., 2010; Schlez et al., 2021; Sing et al., 2016). To our knowledge, strains of *C. belfantii* were only isolated from human respiratory samples.

Despite previous reports of animal Cdc infections *e*.*g*., (Lartigue et al., 2005; Leggett et al., 2010; Hacker et al., 2016a; Abbott et al., 2020), no large case series of such infections, including precise bacterial identification and genotyping, have been reported. Here, we present a case series of Cdc infections in dogs, cats, rabbits, rats and horses in France over 2 years, and report their clinical and microbiological characteristics.

## Materials and methods

### Inclusions and microbiological characterization

A total of 18 308 samples were analyzed during the period August 2019 to August 2021 (24 months), originating from sick animals (dogs, cats, horses and small mammals) from metropolitan France (**Table 1**). All samples from which isolates were identifed as members of the Cdc were included in the present study. Although *C. pseudotuberculosis* was sometimes recorded, this species was excluded because the shipping and reception process of these samples are known to be biased (samples of livestock are generally sent to state laboratories and only occasionally to private laboratories).

**Table 1.** Overview of samples screened and sources with isolates of Corynebacteria of the diphtheriae complex.

| Clinical source | No. Samples screened | *C. ulcerans*, tox+ | *C. ulcerans*, tox- | *C. rouxii* (all tox-) | *C. diphtheriae* (tox+) | Total *Cdc* % |
| --- | --- | --- | --- | --- | --- | --- |
| Dog |  |  |  |  |  |  |
| ear swab | 9439 (52%) | 3 | 6 | 8 | 0 | 17 (0.2%) |
| respiratory tract sample | 1037 (6%) | 1 | 3 | 0 | 0 | 4 (0.4%) |
| wound/skin swab | 2834 (15%) | 7 | 8 | 1 | 0 | 16 (0.6%) |
| Cat |  |  |  |  |  |  |
| ear swab | 1758 (10%) | 1 | 2 | 1 | 0 | 4 (0.2%) |
| respiratory tract sample | 1191 (7%) | 6 | 4 | 0 | 0 | 10 (0.8%) |
| wound/skin swab | 1027 (6%) | 2 | 3 | 1 | 0 | 6 (0.6%) |
|  |  |  | 1 (cystitis) |  |  |  |
| Horse |  |  |  |  |  |  |
| ear swab | 5 (0.03%) | 0 | 0 | 0 | 0 | 0 (0%) |
| respiratory sample | 111 (0.6%) | 0 | 0 | 0 | 0 | 0 (0%) |
| wound/skin swab | 121 (0.7%) | 0 | 0 | 0 | 2 (dermatitis/conjunctivitis) | 2 (1.6%) |
| Rat |  |  |  |  |  |  |
| ear swab | 6 (0.03%) | 0 | 0 | 0 | 0 | 0 (0%) |
| respiratory sample | 20 (0.1%) | 3 | 0 | 0 | 0 | 3 (15%) |
| wound/skin swab | 10 (0.05%) | 0 | 0 | 0 | 0 | 0 (0%) |
| Rabbit |  |  |  |  |  |  |
| ear swab | 116 (0.6%) | 0 | 0 | 0 | 0 | 0 (0%) |
| respiratory sample | 289 (1.6%) | 1 | 0 | 0 | 0 | 1 (0.3%) |
| wound/skin swab | 127 (0.7%) | 0 | 0 | 0 | 0 | 0 (0%) |
| Others (goat, sheep, exotics...) |  |  |  |  |  |  |
| ear swab | 28 (0.2%) | 0 | 0 | 0 | 0 | 0 (0%) |
| respiratory sample | 84 (0.5%) | 0 | 0 | 0 | 0 | 0 (0%) |
| wound/skin swab | 105 (0.6%) | 0 | 0 | 0 | 0 | 0 (0%) |
| **Total** | **18308 (100%)** | **24** | **27** | **11** | **2** | **64 (0.3%)** |

The geographical location of the cases was recorded using the corresponding French administrative Department (in metropolitan France, there are 96 geographic and administrative divisions called “Départements”). Clinical data (age, sex and clinical symptoms) were collected using information communicated on the order form.

Sterile swabs with liquid Amies culture medium were used to sample the affected animals, and were sent to the laboratory under cooled conditions. There, swabs were plated on solid culture media (Columbia agar additioned with 5 % sheep blood and colistine + nalidixic acid (CNA) under an extractor hood. Solid culture media were incubated at 35°C under 5% enriched CO2 atmosphere during 24 to 48h. Isolates belonging to the *Corynebacteria* of the *diphtheriae* complex form dry colonies of dark yellow color on the CNA culture medium. The identity of the suspected colonies was confirmed by Matrix Assisted Laser Desorption Ionization Time of Flight Mass Spectrometry (MALDI-TOF MS, Bruker Daltonics, Germany).

Antibiotic susceptibility testing was performed by disk diffusion on Mueller-Hinton culture medium supplemented with 5% blood following the instructions of the French AFNOR norm NFU-47. Diameters of the inhibition zone were read and interpreted using software tool SIRWeb (i2A, France). As there are no veterinary breakpoints, the 2013 guidelines from the human CASFM (https://www.sfm-microbiologie.org/wp-content/uploads/2020/07/CASFM_2013.pdf) were applied as requested by law. The antibiotic susceptibility testing was composed of commonly used antibiotics in veterinary clinics, including beta-lactams (amoxicillin), tetracyclines (doxycycline and tetracycline), aminoglycosides (gentamicin), macrolides (erythromycin, azithromycin, spiramycin) and others (trimethoprim-sulfamethoxazole). Recent data suggest that *C. ulcerans* is inherently resistant to clindamycin (Marosevic et al., 2020a). Therefore, clindamycin was only tested for *C. diphtheriae* and *C. rouxii*. As there are no breakpoints for veterinary fluoroquinolones and cefovecin, these antibiotics were not tested.

All *Cdc* isolates were sent to the National Reference Center for corynebacteria of the *diphtheriae* complex for confirmation of the identification and for the detection of the diphtheria toxin *tox* gene by real time PCR (Badell et al., 2019). This multiplex assay consists of amplifying a fragment of the *rpoB gene* with primer sets specific for either (i) the species *C. diphtheriae, C. belfantii* or *C. rouxii*; or (ii) species *C. ulcerans* and *C. pseudotuberculosis*; in addition, a fragment of the *tox* gene is detected. The production of the diphtheria toxin was assessed using the modified Elek test (Engler et al., 1997).

### Multilocus sequence typing (MLST)

Isolates were retrieved from −80°C storage and plated on tryptose-casein soy agar for 24 to 48□h. A small amount of bacterial colony biomass was resuspended in a lysis solution (20□mM Tris-HCl [pH 8], 2 mM EDTA, 1.2% Triton X-100, and lysozyme [20□mg/ml]) and incubated at 37°C for 1 hour, and DNA was extracted with the DNeasy Blood&Tissue kit (Qiagen, Courtaboeuf, France) according to the manufacturer’s instructions. MLST was performed as previously described (Bolt et al., 2010; Guglielmini et al., 2021); alleles and profiles were defined using the BIGSdb-Pasteur platform (https://bigsdb.pasteur.fr/diphtheria).

### Statistical analyses

Dogs and cats were divided into 3 groups of age: under 2 years of age; 2 to 8 years; and older than 8 years. A few dogs could not be attributed to a group because the information was not available. Pearson’s chi-squared statistic was used to compare the different characteristics for the categorical variables. Statistical tests were performed in SPSS Statistics software version 25 (IBM, New York, USA).

### Ethical statement

Animals were sampled for diagnostic purpose. No ethics approval was requested for this retrospective study.

## Results

### Isolation of members of the *C. diphtheriae* species complex in animals in France

Between August 2019 and August 2021, 18 308 sick animals (dogs, cats, horses and small mammals) were sampled from veterinary clinics from across metropolitan France. There were 2732 nasal swabs, 4224 cutaneous/wound/abscess swabs and 11 352 auricular swabs (**Table 1**). A total of 64 consecutive, non-duplicated isolates belonging to the Cdc were identified from these samples (**Table 2)**. Of these, 26 were carriers of the *tox* gene coding for the diphtheria toxin, including 24 *C. ulcerans* and 2 *C. diphtheriae*. Of the 38 non-toxigenic Cdc isolates, 27 belonged to *C. ulcerans* and the 11 remaining ones were *C. rouxii*. No non-toxigenic *C. diphtheriae, C. pseudotuberculosis* or *C. belfantii*, were detected.

**Table 2.**
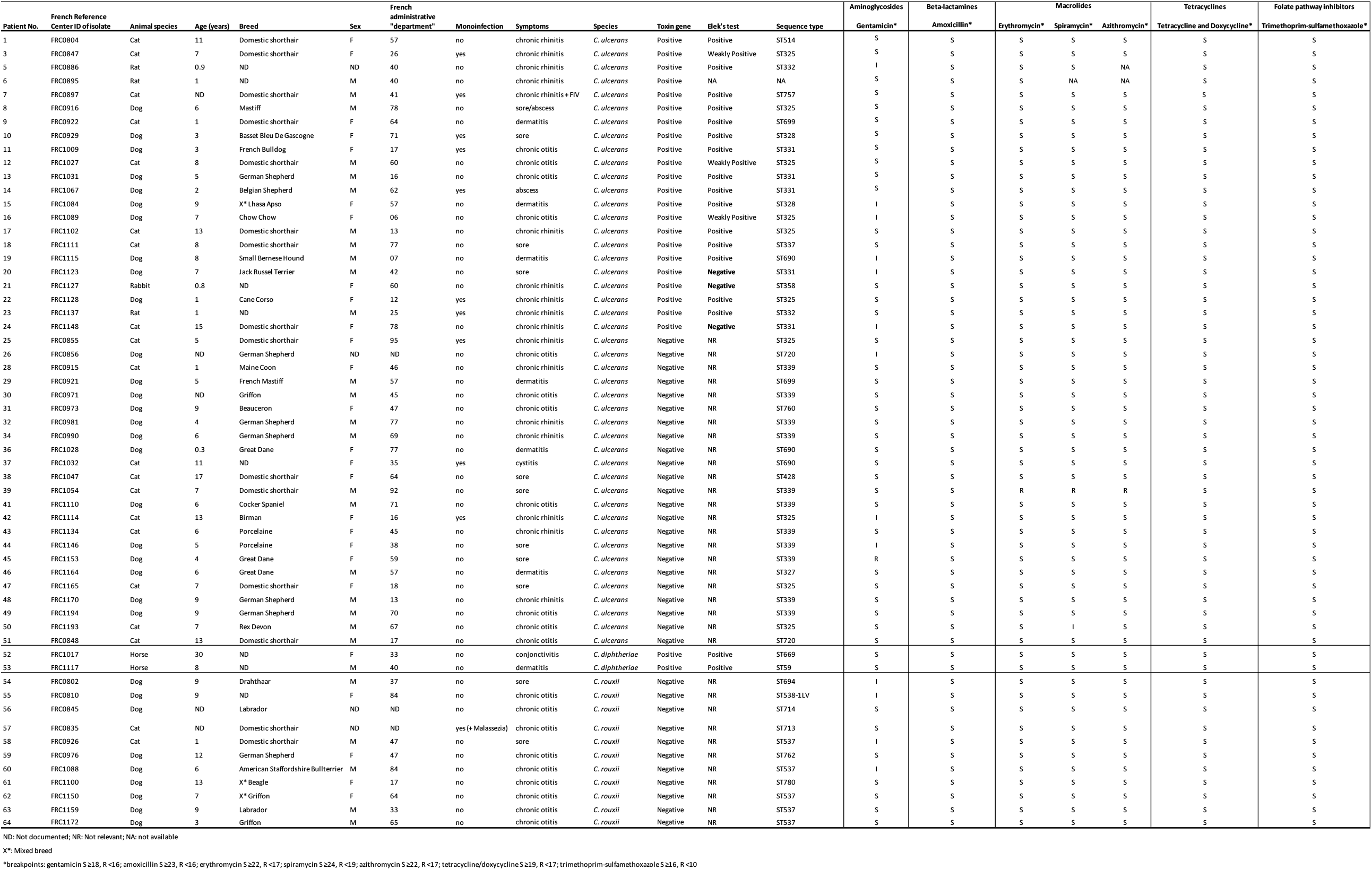
Strain characteristics.

The production of the diphtheria toxin was assessed for *tox*-gene bearing islates. Both toxin gene-carrying *C. diphtheriae* strains were positive in the Elek’s test. Among the 24 *tox*-positive *C. ulcerans*, one strain (FRC0895) was not available for testing, and 20 of the 23 tested isolates (87.0%) had a positive result for Elek’s test, including 3 isolates with a weakly positive Elek test result. These three isolates belonged to a single genetic subtype, ST325. Three *tox*-positive *C. ulcerans* (2 of ST331, 1 of ST358) were negative for the Elek test, hence corresponding to non-toxigenic, *tox* gene-bearing (NTTB) strains.

Of the 64 infections with Cdc isolates, 37 were in dogs, 21 in cats, 3 in rats, 1 in a rabbit and 2 in horses (**Tables 1 and 2**). The age of dogs and cats ranged from 1 year to 13 years of age. There was no statistically significant difference in prevalence of Cdc isolates among the different age groups. The most frequent breed among the *C. ulcerans* dog cases was the German Shepherd (9 of 28 dogs of this breed). Dogs belonging to this breed were strongly associated with infections caused by *C. ulcerans:* there were 556 German Shepherd out of 13310 dogs; p < 0.00001.

### Antimicrobial susceptibility of Cdc isolates

*C. ulcerans* isolates were susceptible to most antibiotics (**Table 2**). Spiramycin was tested in 50 isolates, following the the publication of Abott et al. in 2020 (Abbott et al., 2020); all were susceptible, except one resistant and one susceptible at higher concentration (previously ‘intermediate’). Azithromycin was tested in 49 samples, and only one isolate (*C. ulcerans* FRC1054) was resistant; this isolate was also resistant to erythromycin and spiramycin. Nine *C. ulcerans* isolates were tested ‘intermediate’ for gentamicin, and one was resistant, whereas the 42 other ones were susceptible. The *C. ulcerans* isolates were also susceptible to erythromycin in all but one case, and were all susceptible to tetracyclins and trimethoprim-sulfamethoxazole.

The two *C. diphtheriae* isolates were susceptible to all antibiotics tested, including clindamycin. The *C. rouxii* isolates were susceptible to amoxicillin, tetracyclins, trimethoprim-sulfamethoxazole, erythromycin, spiramycin, azithromycin, clindamycin, but only 7 isolates were susceptible to gentamicin, the 4 others being ‘intermediate’.

### Genotyping of isolates using multilocus sequence typing

MLST analysis showed a wide diversity of *C. ulcerans* isolates, with 15 distinct sequence types (ST; **Figure 2**; **Table 2**). *C. rouxii* was also genetically heterogeneous, with 7 STs, and the two *C. diphtheriae* belonged to ST59 and ST699 (**Figure 2**). Six *C. ulcerans* STs and one *C. rouxii* ST comprised more than one isolate. For these STs, geographical provenance and animal source were heterogeneous, indicating spread across localities and host species (**Figure S1**). Two *C. ulcerans* STs (ST325 and ST690) comprised both *tox*-positive and *tox*-negative isolates (**Figure 2**), implying that the loss or gain of the *tox* gene occurred within these lineages. We noted that *C. rouxii* isolates were originating mainly from South-Western France (**Figures 1 and S1**).

**Figure 1.**
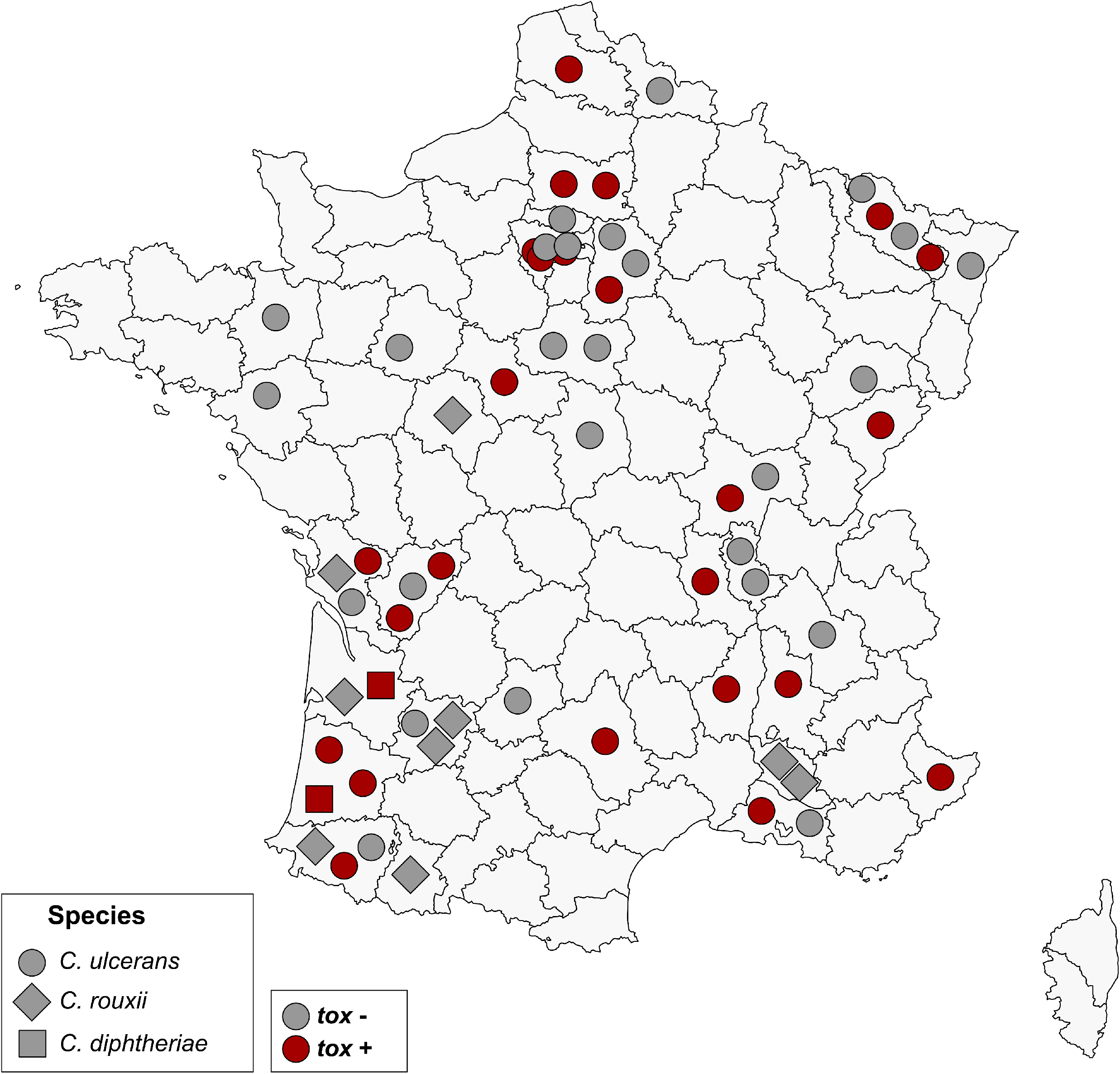
Geographic distribution of Cdc infection cases in pets. The 51 *Corynebacterium ulcerans*, 2 *C. diphtheriae* and 9 *C. rouxii* are displayed within their administrative Departement of origin, with a species-specific symbol. *tox* gene-bearing strains are indicated in red, non-*tox* gene-bearing in grey. For two cases, location was not available and they are therefore not represented here.

**Figure 2.**
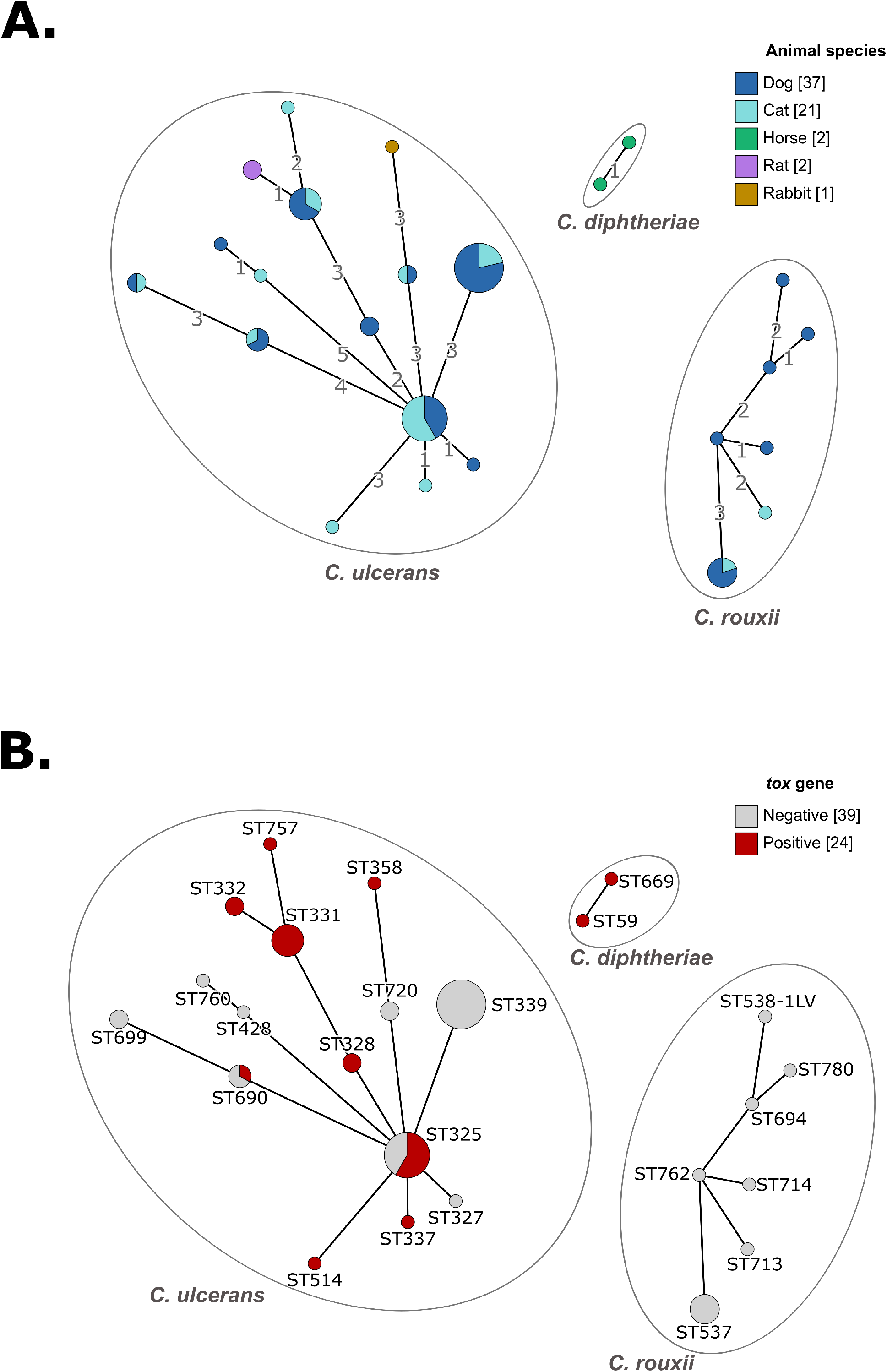
MLST diversity of Cdc isolates from pets. (A) Minimum spanning tree of 7-gene MLST profiles, colored in function of animal host species. (B) Same, colored by diphtheria toxin gene presence. The graphs were obtained using the GrapeTree tool, which is pluged onto the BIGSdb platform (https://bigsdb.pasteur.fr/cgi-bin/bigsdb/bigsdb.pl?db=pubmlst_diphtheria_isolates&page=plugin&name=GrapeTree).

### Clinical characteristics of toxigenic *C. ulcerans* isolates

Eleven of the 24 toxigenic *C. ulcerans* were isolated from nasal swabs. They were taken from 6 cats with chronic rhinitis and 3 rats, 1 dog and 1 rabbit. Two of the rats were bought from the same pet shop, and they were both coinfected with *Staphylococcus aureus*. Euthanasia was chosen because of the zoonotic potential. Four other nasal swabs also harboured microorganisms other than *C. ulcerans*, including *Pasteurella multocida, Escherichia coli, Bacteroides spp, Stenotrophomonas maltophilia* and *Staphylococcus pseudointermedius*.

The second most frequent isolation source was skin (9 of 24 cases) and included wounds, pyoderma and abscesses. Seven of these *C. ulcerans* were sampled from dogs and two from cats. In 4 of the 9 skin cases, *C. ulcerans* was the only microorganism recovered from the samples, while in 5 cases there was a coinfection; organisms encountered were *Pasteurella multocida, Pasteurella canis, Proteus mirabili*s, *Staphylococcus aureus, Staphylococcus pseudointermedius, Streptococcus canis, Pseudomonas aeruginosa* and *Streptococcus canis*.

Last, there were four ear infections by toxigenic *C. ulcerans*. These were observed in three dogs and one cat. In 3 cases, otitis was also associated with at least another microorganism (*Proteus mirabilis, Pseudomonas aeruginosa, Streptococcus cani or Streptococcus dysgalactiae*).

Three of the 24 animals infected by a toxigenic *C. ulcerans* had a bacteriological follow-up examination. Two cats (patients 3 and 4, **Table 2**), had rhinitis with *C. ulcerans* in pure culture, and tested positive again at control screenings 1 month and 18 days after the initial visit, respectively. Patient 3 had been treated 10 days with amoxicillin-clavulanate without success. No further control was performed. In patient 4, a four-week cure of cefovecin did not eliminate the toxigenic *C. ulcerans*. However, a 1-month treatment with amoxicillin-clavulanate, to which the isolate was susceptible, was followed by a negative result for *C. ulcerans* on the control sample 4 months later.

One rabbit (patient 21) with *C. ulcerans* and *Pasteurella multocida* co-infection was still positive for *C. ulcerans* and *P. multocida* despite treatment with marbofloxacin for 7 days and then trimethoprim-sulfamethoxazole for 10 more days. No further control was performed.

### Clinical characteristics of non-toxigenic *C. ulcerans* isolates cases

The 27 non-toxigenic *C. ulcerans* were retrieved from 6 cases of dermatitis (6 dogs), 7 cases of rhinitis (4 cats, 3 dogs), 8 cases of otitis (6 dogs, 2 cats), 1 case of cystitis (cat) et 5 cases of non-healing purulent wounds (3 cats, 2 dogs). In total, there were 10 cats and 17 dogs. Three monoinfections were observed and concerned two cats with chronic rhinitis and one cat with cystitis.

The other infections were polymicrobial. Regarding the otitis cases, the polyinfections consisted of co-infection with *Streptococcus canis* (in 3 cases), *Pseudomonas aeruginosa* (n=2), *Proteus mirabilis* (n=2), *Staphylococcus pseudointermedius* (n=2) and *Klebsiella pneumoniae* (n=1). The polyinfections in the cases of rhinitis were co-infections with *Staphylococcus intermedius* (n=2), *Pasteurella multocida* and *Bordetella bronchiseptica* (n=1), *Klebsiella oxytoca* (n=1) and *Staphylococcus aureus* (n=1). The dermatitis polyinfections were co-infections with *S. pseudointermedius* (n=5), *P. mirabilis* (n=3) and *Pasteurella canis* (n=1). Last, polyinfections of sores comprised *Escherichia coli* (n=2), *S. aureus/pseudointermedius* (1 case each), *Pasteurella dagmatis/multocida* (1 case each) and *Streptococcus dysgalactiae* (n=1).

There was only one follow up available (patient 25). It was a 5 year old cat with chronic rhinitis and a *C. ulcerans* monoinfection. This patient remained positive for non-toxigenic *C. ulcerans* during 6 months. In between, the cat was treated with doxycycline to which the strain was tested susceptible in the 2 antibiotic susceptibility testings before and after treatment.

### Clinical characteristics of *C. diphtheriae* and *C. rouxii* isolates cases

The two cases of *C. diphtheriae* were toxigenic and consisted of conjonctivitis and pastern dermatitis in two horses. Both cases were polymicrobial infections, in which *S. dysgalactiae* (conjonctivitis) and *S. aureus* and *Enterobacter cloacae* (pastern dermatitis) were also found.

*C. rouxii* infections were observed in 9 dogs and 2 cats and were all non-toxigenic. It consisted of were 9 otitis cases (8 dogs and 1 cat) and two remaining non-healing wounds (one dog and one cat). *C. rouxii* was identified retrospectively by MLST, as this species was only recognized as a separate species during the study period (in 2020), and as MALDI-TOF still identifies *C. rouxii* as *C. diphtheriae*, based on the MALDI Biotyper Compass database version 4.1 (version (100).

Almost all *C. rouxii* infections were polymicrobial infections. There was only one mono-bacterial infection (ear infection of a cat), though *Malassezia* spp. was also identified (**Table 2**). The most frequent (n=5) co-infecting bacteria was *P. aeruginosa* as expected in case of otitis. Other co-infecting bacteria were *P. mirabilis* (n=3), *Staphylococcus sp*. (n=2), *S. canis* (n=2), *Corynebacterium amycolatum* (n=1), *Citrobacter koseri* (n=1) and anaerobic bacteria (n=1). The age of the infected animals here ranged from 1 year to 13 years of age with no statistically significant difference between age groups.

## Discussion

We report on the occurrence, clinical and microbiological characteristics of infections caused in companion animals by members of the Corynebacteria of the *diphtheriae* complex in France. Although *C. ulcerans* was the most frequent species, *C. rouxii* and, more rarely, *C. diphtheriae* were also found.

All isolates initially identified as *C. diphtheriae* from dogs and cats were in fact *C. rouxii*, as identified by MLST gene sequences. This important observation suggests that this novel species has been overlooked, and in fact appears more prevalent than *C. diphtheriae* itself in animal infections. As all *C. rouxii* infections, mostly of ears, were co-infections with other bacteria or the yeast *Malassezia*, this species may represent a commensal in dogs and cats. So far, all *C. rouxii* isolates are *tox*-negative, with the exception of isolates from two cats from the USA (Hall et al., 2010) but in these two isolates, the *tox* gene was disrupted and therefore the toxin was not produced. Previously, *C. rouxii* has been isolated from cutaneous infections, vascularitis and peritonitis in humans and a purulent orbital cellulitis in a dog (Badell et al., 2020). Our study correspond to the largest group of infections with *C. rouxii* reported so far. We suggest that *C. rouxii* may represent a novel zoonotic pathogen, as pets may clearly serve as a reservoir for these human infections. However, so far, no case of transmission of *C. rouxii* between human and animals was documented. *C. rouxii* was more frequently isolated in South-Western France (**Figure 1**), and was genetically diverse (**Figure 2 and Figure S1**). Although the sampling is still limited, this may reflect the existence of local conditions that favor the infections of dogs or cats by *C. rouxii* in this part of France.

Toxigenic *C. diphtheriae* was isolated from 2 horses. Mixed wound infections in horses were previously described (Henricson et al., 2000; Leggett et al., 2010), and colonisation with *C. diphtheriae* was reported in 6.9% of slaughter horses in Romania (Stănică et al., 1968). The pathogenic potential of *C. diphtheriae* in horses is questionable given its report from polymicrobial infections or asymptomatic carriage. Given the well established human-to-human transmission of *C. diphtheriae*, and possible asymptomatic carriage, the possibility should be considered that the detection of this pathogen in horses corresponds to reverse zoonosis.

*C. ulcerans* is now well-established as a zoonotic member of the Cdc. Whereas no human-to-human transmission was reported, human cases have often been associated with animal contacts, and in several cases the genetic fingerprinting of isolates supported an epidemiological link between the animal and human isolates, strongly establishing the zoonotic character of *C. ulcerans* (Lartigue et al., 2005; Othieno et al., 2019; Vandentorren et al., 2014). The emergence of *C. ulcerans* has been noted in France, Germany and the UK (Haut Conseil de la santé publique, 2019; Robert Koch-Institut, 2018; Wagner et al., 2010).

This is the first study based on the systematic analysis of a very large animal cohort (18 308 samples), which provides data on the frequency of *C. ulcerans* in various types of clinical samples from animals. Abbott and colleagues (Abbott et al., 2020) found 7 *C. ulcerans* among 804 nasal samples (0.87%): 3 samples from 668 dogs, and 4 samples from 64 cats, wheras Katsukawa et al. (Katsukawa et al., 2012) found 44 *C. ulcerans* in 583 pharyngeal samples of dogs (7.5%). Here, we found 51 *C. ulcerans*, including 24 *tox*-positive ones, in 18 308 clinical samples (0.27%), mostly in nasal swabs (18 *tox*-positive and negative samples).

Given the wide geographic distribution of cases (**Figures 1 and S1**), the reporting of *C. ulcerans* in this study does not seem affected by a bias caused by local events of transmission. In addition, the genetic diversity of *C. ulcerans* isolates indicates that the reporting of this pathogen cannot be attributed to the clonal spread of a single emerging strain of *C. ulcerans*. Whether the increase in cases observed over the two last decades is due to changes in diagnostic practice (*i*.*e*., MALDI-TOF) or increasing awareness of the necessity to report and test for the presence of the diphtheria toxin, rather than a real epidemiological phenomenon of emergence, remains unclear. Before 2019, *C. ulcerans* of animal origin were only sent sporadically to the national reference laboratory. Awareness of this zoonotic bacteria remains low among veterinarians and veterinary laboratories, among which it is often considered a commensal bacteria of animals.

*C. ulcerans* is widely distributed, having been reported from multiple animal species (Hacker et al., 2016a). Whether pet animals represent a natural reservoir, or (more probably) are themselves contaminated by *C. ulcerans* from other animal species or sources, is an important question. Toxigenic *C. ulcerans* infections were reported in wildlife carnivores such as foxes, otters and owls (Foster et al., 2002) and insectivores (hedgehogs and Japanese shrew owls) (Katsukawa et al., 2016b), whereas non-toxigenic *C. ulcerans* infections seem to be more frequent in omnivores (wild boars) and herbivores (roe deer) (Berger et al., 2019a). Only wild animals with major symptoms are diagnosed, while the asymptomatic carriership is rarely investigated (Berger et al., 2019b). Transmission by predatory hunting was supported by a study showing high serum diphtheria antitoxin titers in hunting dogs (Katsukawa et al., 2016a). A reservoir of symptomatic or asymptomatic carriers in small wild mammals such as herbivores, lagomorphs or rodents, which may be preys to dogs and cats, should be further investigated. Here, we reported 4 cases of toxigenic *C. ulcerans* in rats or rabbits. Although the cases described in this study are from symptomatic animals, an asymptomatic carrier state of *C. ulcerans* in dogs, cats, horses, rodents and wild life animals has been described in other studies (Dias et al., 2010; Katsukawa et al., 2016a, 2016b; Othieno et al., 2019; Vandentorren et al., 2014). Future studies should investigate the presence of Cdc in healthy animals to better understand and control its transmission to pets.

There is evidence for a pathogenic potential of *C. ulcerans* in animals (Hacker et al., 2016a). Here, *C. ulcerans* was often isolated from nasal swabs in cats with chronic rhinitis. While acute feline upper respiratory infections are mostly caused by viruses (*Feline Herpesvirus-1* and *feline Calicivirus*) (Lappin et al., 2017) and are most of the time self-limiting, chronic rhinitis is more concerning. In general, bacteriological examinations performed on nasal swabs are often not useful because of the presence of commensal flora and only a few bacteria are considered primary pathogens (Lappin et al., 2017). Toxigenic *C. ulcerans* may be implicated in the clinical manifestation of rhinitis. Importantly in 7 cases of rhinitis, *C. ulcerans* was the only infective agent retrieved from the nasal swabs, a fact that strongly suggests a primary pathogenic nature of *C. ulcerans* in cats, consistent with a previous study (Abbott et al., 2020). A limitation of the present study is that it was perfomed retrospectively; hence, the final diagnosis as well as the complete clinical picture were not available.

The pathogenicity of non-toxigenic strains is not well understood. Here, a few monoinfections by non-toxigenic *C. ulcerans* (2 cases of rhinitis and 1 cystitis) are reported. The infections could be favoured by other virulence factors of *C. ulcerans*, such as phospholipase D (Hacker et al., 2016a).

Large dog breeds were disproportionally affected by *C. ulcerans*, and 32% of them were German Shepherds. This interesting observation might be explained by a closer contact with small mammals, as large dogs are generally kept outdoors. But it could also reflect a breed predisposition, as German Shepherd dogs are more susceptible to pyoderma (Rosser, 2006) and other bacterial infections, for example by *Ehrlichia canis* (Nyindo et al., 1980). Six cases of German Shepherd in this study presented with pyoderma or otitis. Further studies will be needed to investigate a possible dog breed predisposition to *C. ulcerans*.

In at least 3 cases of toxigenic *C. ulcerans* and 1 case of non-toxigenic *C. ulcerans* (patient numbers 3, 4, 21 and 25), the infection persisted, despite antibiotic susceptibility, as reported in previous works (Lartigue et al., 2005; Vandentorren et al., 2014; Abbott et al., 2020). One speculative possibility could be that chronic infections might involve some intracellular bacteria (Moore et al., 2015), which would be more accessible to macrolides, which can reach higher intracellular concentrations compared to beta-lactams. Systemic infections by *C. diphtheriae* suggest that this pathogen is not only able to attach to host epithelial cells, but is able to gain access to deeper tissues and to live intracellulary (Ott et al., 2010). *Corynebacterium* guidelines in humans recommend the use of beta-lactams or macrolides. Yet, while chronic human carriers can be treated with azithromycin or rifampicin, the use of the latter is forbidden in the veterinary field.

Recent data suggest to use spiramycin in cases of animal infection by *C. ulcerans*. A 10-day course of the combination of spiramycin and metronidazole was successful to clear *C. ulcerans* from a dog, and a 6-day course of the same antibiotic combination was successful in cats (Abbott et al., 2020; UK Health Security Agency, 2022). As spiramycin only exists in a combination with metronidazole, and as metronidazole has side effects (neurotoxicity and effect on the microbiote), azithromycin seems to be a better choice for treatment.

The efficiency of the antibiotic treatment should be verified ideally by 2 samples, taken 5 to 7 days following completion of the antibiotic course. The macrolide agent (azithromycin rather than a spiramycin/metronidazole combination) and the optimal treatment duration should be investigated by future studies.

This work is relevant to the development of veterinarian guidelines in case of Cdc infections in animals. Currently, in France no recommendations are established around the detection of toxigenic Cdc in animals, either for their animal or human contacts. From a public health perspective, the zoonotic potential of toxigenic *C. diphtheriae* or *C. ulcerans* in chronic skin infections, non-healing wounds, rhinitis and otitis, justifies investigating these body locations for such pathogens, and these samples should not be considered to reveal only commensal organisms. This would be more in line with the fact that in case of diphtheria due to toxigenic *C. ulcerans* in a human, recommendations exist in France that contact animals should be sampled and treated (Haut Conseil de la santé publique, 2011, 2019, 2021).

Toxigenic corynebacteria in animals are not mandatory to notify, and the costs of treatment, and clearance and sampling of contact animals, clearly represent a limitation for the implementation of follow-up investigations and control measures. Asymptomatic human carriers of toxigenic strains are treated with the same antibiotic regimen as symptomatic cases, with clearance swabs to ensure eradication. From an epidemiological viewpoint, it would be coherent to follow the same practice for animals, and this should be discussed between veterinary and human health institutions.

## Acknowledgements

We thank Stéphanie Gilles from the bacteriology department of Cerba Vet for her support, and Annie Landier and Nathalie Armatys (Institut Pasteur) for technical help.

## Authors license statement

This research was funded, in whole or in part, by Institut Pasteur and by European Union’s Horizon 2020 research and innovation programme. For the purpose of open access, the authors have applied a CC-BY public copyright license to any Author Manuscript version arising from this submission.

## Ethical approval statement

Authors declare ethics approval was not needed for this retrospective study.

## Author contributions

KM and SB conceived, designed, and coordinated the study. GR and EB performed the microbiological cultures of the isolates and their biochemical and molecular characterizations. MH analyzed the sequence data and created the figures. GA and KM reviewed the clinical source data of the isolates. KM and SB wrote the initial version of the manuscript. All authors provided input to, read, and approved the final manuscript.

