## Supplementary material for "Corynebacterium of the *diphtheriae* complex in companion animals: clinical and microbiological characterization of 64 cases from France"

**Supplementary Figure S1**. **Geographic distribution of the MLST sequence types (ST).**

Colors correspond to STs, as indicated on the Minimum spanning tree and on the key. For two cases, location was not available and there are therefore not represented here.

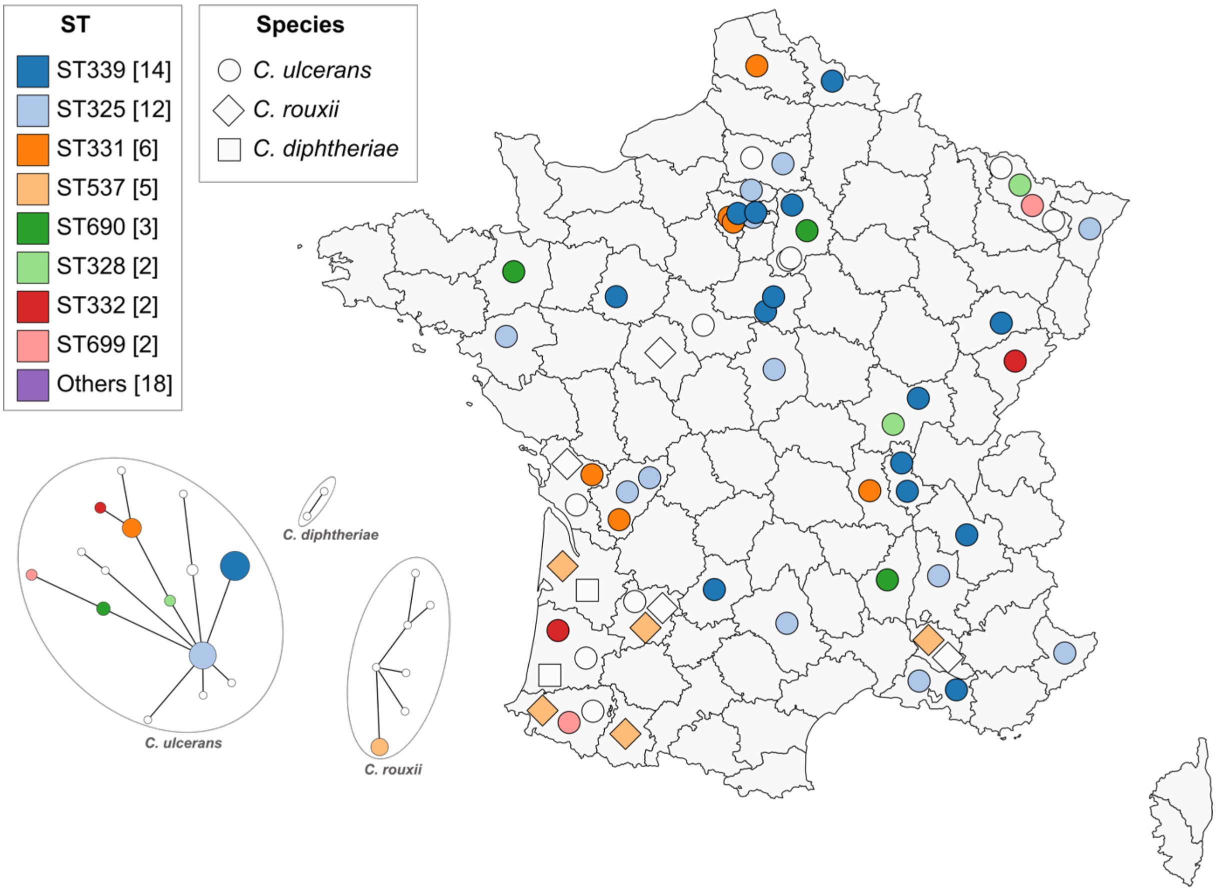
